# Compensatory mutations are associated with increased *in vitro* growth in resistant clinical samples of *Mycobacterium tuberculosis*

**DOI:** 10.1101/2023.06.21.545231

**Authors:** Viktoria Brunner, Philip W Fowler

**Affiliations:** Nuffield Department of Medicine, University of Oxford, Oxford, UK; National Institute of Health Research Oxford Biomedical Research Centre, John Radcliffe Hospital, Headley Way, Oxford, UK; Health Protection Research Unit in Healthcare Associated Infections and Antimicrobial Resistance, University of Oxford, UK

## Abstract

Mutations in *Mycobacterium tuberculosis* associated with resistance to antibiotics often come with a fitness cost for the bacteria. Resistance to the first-line drug rifampicin leads to lower competitive fitness of *M. tuberculosis* populations when compared to susceptible populations. This fitness cost, introduced by resistance mutations in the RNA polymerase, can be alleviated by compensatory mutations (CMs) in other regions of the affected protein. CMs are of particular interest clinically since they could lock in resistance mutations, encouraging the spread of resistant strains worldwide. Here, we report the statistical inference of a comprehensive set of CMs in the RNA polymerase of *M. tuberculosis*, using over 70,000 *M. tuberculosis* genomes that were collated as part of the CRyPTIC project. The unprecedented size of this data set gave the statistical tests to investigate the association of putative CMs with resistance-conferring mutations much more power. Overall, we propose 51 high-confidence CMs by means of statistical association testing and suggest hypotheses for how they exert their compensatory mechanism by mapping them onto the protein structure. In addition, we were able to show an association of CMs with higher *in vitro* growth densities, and hence presumably with higher fitness, in resistant samples in the more virulent *M. tuberculosis* Lineages 2 and 3. In Lineage 2, our results even suggest the association of CM presence with significantly higher *in vitro* growth than for wild-type samples, although this association is confounded with lineage and sub-lineage affiliation. Our findings emphasise the integral role of CMs and lineage affiliation in resistance spread and increases the urgency for antibiotic stewardship, which implies accurate, cheap and widely accessible diagnostics for *M. tuberculosis* infections to not only improve patient outcomes but also to prevent the spread of resistant strains.

## Introduction

The rise of multidrug-resistant bacteria is one of the grand challenges we are facing as a global society. Almost a century ago, the fight against bacterial infectious diseases seemed to be near its end due to the discovery of potent antibiotics like penicillin. ^1;2^ Yet, a study in 2019 found 7.7 million deaths per year associated with bacterial infections. ^3^ This ongoing crisis is mainly caused by the short generation times, high mutation rates and genetic recombination of pathogenic bacteria which make it possible for antimicrobial resistance to arise within short time frames and spread rapidly throughout bacterial populations. ^4^ Modern societies contribute to the problem by inappropriately administrating antibiotics, encouraging the faster emergence and spread of multidrug-resistant strains through directional selection. ^2^ New antibiotic drugs are being developed but the pace is slow and the rate at which resistance to new drugs develops is higher. ^5^ With the increasing prevalence of antibiotic resistance and the decreasing discovery rate of potent new drugs, we are heading towards a human-fabricated global health crisis. ^4;6^

*M. tuberculosis* is the etiological agent of tuberculosis (TB) and prone to developing resistance to major antibiotics. ^7^ Hence, although the first antibiotic for treatment of TB was identified as early as 1948, ^8^ the disease remains responsible for the death of about 1.6 million people per year. In addition, the proportion of people infected with *M. tuberculosis* strains resistant to the major first-line antibiotic drugs is rising. ^9–11^ Understanding how resistance mutations emerge, how they become fixed and how they spread is hence of high importance.

One of the four first-line antibiotics for treating TB is the drug rifampicin (RIF) which binds to the RNA polymerase (RNAP). The RNAP holoenzyme in *M. tuberculosis* is comprised of five subunits (*α*_2_*ββ^′^γ*) and an additional *σ* factor (Figure 1A). ^12^ RIF binds close to the active site in the *β* subunit (*rpoB* gene). In susceptible bacteria, amino acids within the so-called “rifampicin resistance determining region” (RRDR) form hydrogen bonds with RIF (Figure 1B). The bound RIF sterically obstructs the elongation of newly synthesized RNA, thereby stalling protein production in the bacterium. ^13^ Since *M. tuberculosis* exhibits very little evidence of horizontal gene transfer, ^14^ resistance to this drug mostly arises through chromosomal mutations within or close to the RRDR, that prevent RIF from binding. ^15–17^ These resistance-conferring mutations introduce different amino acid side chains close to the active site of the RNAP, hence it is not surprising that they also introduce a fitness cost. ^18;19^ This fitness cost in RIF-resistant bacteria manifests clearly in decreased performance in competition assays with susceptible bacteria. ^19^ While the molecular basis of the fitness deficit is not entirely understood, it is suspected to be related to decreased stability of the RNAP open-promoter complexes ^20^ as well as steric hindrance of RNA exit by the mutated amino acids. ^21^ The low fitness phenotype has been observed to be partially or completely rescued by the presence of so-called compensatory mutations (CMs). ^22–24^ These CMs emerge in various subunits of the RNAP ^23^ and have e.g. been shown to partially restore RNAP activity in *Mycobacterium smegmatis*. ^24^ The existence of CMs gives a plausible explanation for the persistence of resistance mutations long after antibiotic treatment is stopped, when the fitness cost should lead to reversion to the wild-type phenotype.

**Figure 1:**
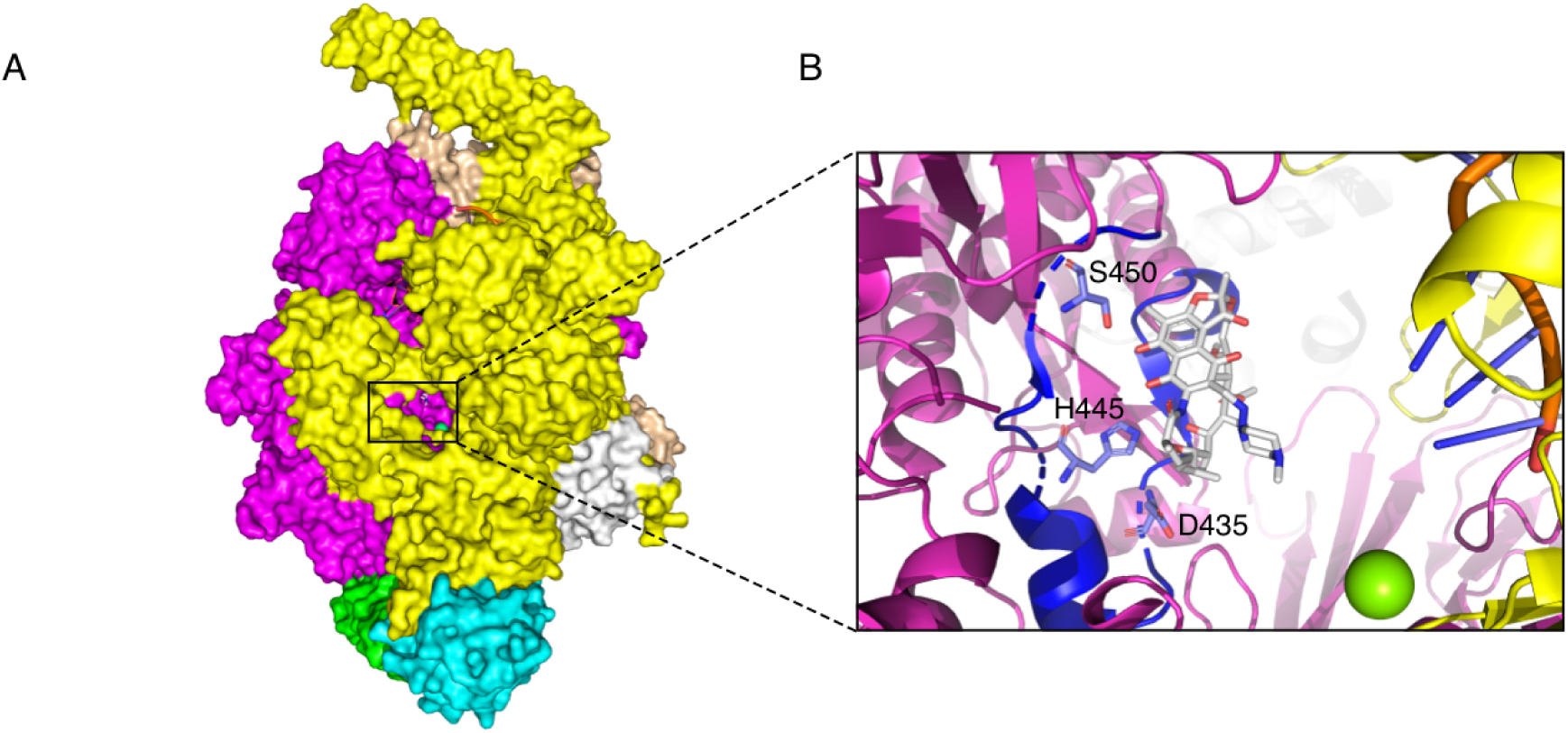
The drug rifampicin (RIF) interrupts RNA synthesis by binding to the *β* subunit of the RNA polymerase (RNAP) ^13^: **(A)** Overview of the entire RNAP. The *β* subunit of the RNAP is shown in magenta, the *β ^′^* subunit in yellow, the two *α* subunits in light blue and green, the *ω* subunit in white and the *σ* factor in light orange. The active site is framed in black. It can be seen through the secondary channel, with the active site magnesium depicted in green and the drug RIF in white. **(B)** Close-up of the active site. The DNA strand used as a template for transcription is shown on the right in orange and dark blue. The *β* subunit of the RNAP is shown in magenta, with the ‘rifampicin resistance determining region’ (RRDR) highlighted in dark blue. The protruding amino acids D435, H445 and S450 are reported to form hydrogen bonds or van der Waals interactions with the drug RIF. ^13^ Due to the proximity of the RRDR to the RNAP active center, the binding of RIF causes a disruption of the RNA synthesis due to steric clash.

Because of their potential to lead to fixation of resistance mutations ^21^ and the attendant epidemiological consequences, CMs have been extensively investigated. Many candidates have been identified in previous studies, ^25–31^ but the number of samples available to these studies is often small. Furthermore, the effects on growth and polymerase activity have only been experimentally confirmed for a small subset of putative CMs. ^24;30^ To further our understanding of resistance spread and persistence, it would be useful to construct a more comprehensive list of CMs and, if possible, to dissect their direct influence on the fitness of *M. tuberculosis*.

The identification of CMs is not trivial and depends on how they are defined. In some publications, due to the low number of available samples, CMs are simply assumed to be all mutations in the RNAP that co-occur with resistance mutations. ^27^ Higher numbers of samples allow CMs to be defined as mutations that *exclusively* occur with resistance mutations. ^25;29;31^ With the dataset of 77,860 *M. tuberculosis* sample genomes we have at hand, where sequencing errors and sample mislabeling are likely to lead to a considerable amount of false positives, it is necessary to employ statistical association testing.

In this paper, we identify a comprehensive set of 51 putative CMs based on a sequenced collection of 77,860 *M. tuberculosis* sample genomes ^32^ collated by the international CRyPTIC project. ^32^ The large size of the dataset ensures the statistical analysis is about 10-100 times more powerful than previous studies. ^25–31^ Another advantage of our investigation is that we have *in vitro* growth data available for a subset of 15,211 samples derived from photographs of 96-well plates after two weeks incubation. ^33^ This allows us to correlate observed growth phenotypes with the respective genotype, i.e. the presence of resistance-conferring and compensatory mutations.

## Results

### Rifampicin resistant *M. tuberculosis* samples show lower *in vitro* growth densities than pan-susceptible samples

First, we analysed the effect of resistance-conferring mutations on the *in vitro* growth of *M. tuberculosis* bacteria. Based on our dataset of 15,211 whole genome sequenced patient-derived *M. tuberculosis* samples with associated 96-well growth data, we defined two sample sets: one resistant to rifampicin (RIF) and another that is susceptible. Resistance was defined as the presence of any mutation associated with resistance to RIF according to a published catalogue, ^34^ whilst a sample was assumed to be susceptible if the catalogue classified it as pan-susceptible (susceptible to all four first line tuberculosis drugs – RIF, isoniazid, ethambutol and pyrazinamide). We excluded samples containing any other non-synonymous mutations in the RNA polymerase (RNAP), since they could have a secondary effect on growth. That being said, we cannot exclude an effect due to mutations in other genes. We shall further assume that synonymous mutations in the RNAP have no effect on the growth phenotype, and hence need not be excluded. The resulting sample sizes can be seen in Table 1.

**Table 1:** Median growth of samples with and without indicated resistance mutations. The confidence interval (CI) for the median is calculated using bootstrapping where ‘CI low’ indicates the lower threshold and ‘CI high’ the upper threshold. Mann-Whitney p-value is calculated in reference to pan-susceptible sample growth and n indicates the sample size.

| mutation | median growth [%] | CI low | CI high | p-value<sub>s</sub> | n |
| --- | --- | --- | --- | --- | --- |
| pan-susceptible | 22.1 | 21.6 | 22.7 |  | 5283 |
| any resistance | 17.6 | 16.6 | 18.5 | 2.95e-12 | 795 |
| specific resistance mutation: |  |  |  |  |  |
| *rpoB* S450L | 16.7 | 15.1 | 18.9 | 4.37e-03 | 196 |
| *rpoB* H445Y | 15.5 | 12.0 | 19.2 | 1.26e-02 | 42 |
| *rpoB* D435V | 16.4 | 15.5 | 17.2 | 3.92e-14 | 325 |

There is a significant difference between the growth distributions of our sets (Figure 2A), as confirmed by a significant Mann-Whitney p-value of 2.95e-12 (Table 1). The median growth density in resistant samples was significantly lower than in pan-susceptible samples, hence we conclude that we see higher growth in non-resistant bacteria.

**Figure 2:**
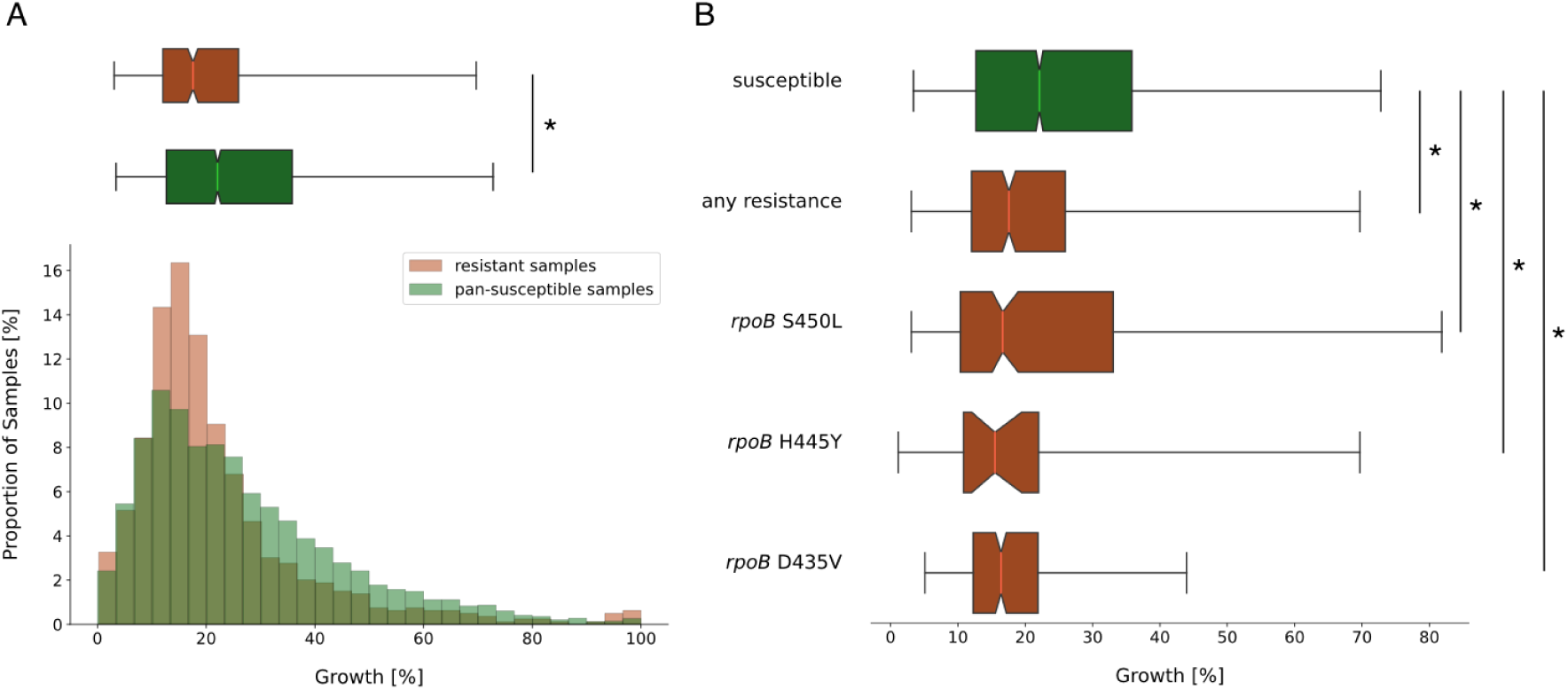
Presence of rifampicin (RIF) resistance-conferring mutations in the RNA polymerase (RNAP) of *M. tuberculosis* is associated with lower median growth compared to pan-susceptible samples. **(A)** Distributions of growth in percent of covered well-area as measured in the CRyPTIC project ^33^ were plotted as a histogram against the proportion of samples that display this amount of growth (bottom) and as a notched box plot reflecting the distribution quantiles (top). Samples with RIF resistance mutations but no other potentially interfering mutations are plotted in red, samples that were classified as pan-susceptible are plotted in green. For the box plot, half of the data lies within the area of the box and 95% in the area covered by the whiskers. Outliers (5 % of the data) were removed to achieve a cleaner representation. Indented areas close to the medians indicate their respective confidence intervals, while the star (*) indicates a significant Mann-Whitney p-value (p *<* 0.05 %). The respective medians, confidence intervals and the Mann-Whitney p-value are listed in Table 1. **(B)** Plot structure equivalent to the box plot in A, but the red bars represent subsets of RIF resistant samples that exhibit only the resistance mutation indicated to their left and no other potentially interfering mutations. The medians, their confidence intervals (CI) and Mann-Whitney p-values of the distributions are listed in Table 1. For a histogram representation of the data please refer to Supplementary Figure S1.

This agrees with work by Gagneux *et al.* who reported that RIF resistance introduces a fitness cost in *M. tuberculosis*, which they show by *in vitro* competitive fitness experiments. ^19^ Their study indicates that the magnitude of the fitness cost in *M. tuberculosis* depends on the mutation, with *rpoB* S450L, the most prevalent RIF resistance mutation observed clinically, being associated with the lowest fitness cost. ^19^ We had sufficient samples to investigate growth for the three most common resistance mutations in the dataset: S450L, H445Y and D435V. Consistent with Gagneux *et al.*, the median growth of *rpoB* S450L mutants was the highest among the three resistance mutations, while still growing to significantly lower densities than pan-susceptible samples (p = 4.37e-03, Figure 2B, Table 1).

### The causal relationship of compensatory mutations with resistance allows identifying compensatory mutations through homoplasy

Since there is a significant relative fitness cost in samples with resistance-conferring mutations and no other mutations in the RNAP, we expected to observe CMs that partially restore fitness in a large proportion of the remaining resistant samples with additional mutations. We performed a Fisher’s exact test for each pair of resistance and co-occurring mutations in the RNAP, to determine if the latter is significantly associated with resistance (Methods).

For a largely clonal species like *M. tuberculosis*, linkage disequilibrium (LD) will artificially inflate the p-values in any SNP-phenotype association test, as can be seen in most genome wide association studies of microbial species. ^35^ LD can hence mask true causal mutation-phenotype relationships, like the one between RIF resistance and CM presence. Furthermore, classic stratification corrections alone will not be sufficient to identify causal variants in species with strong LD and the use of homoplasy as a criterion has been proposed instead. ^36^ We hence decided to apply a very conservative, heuristic p-value cut-off to counter the p-value inflation, and additionally excluded any hits that are not homoplastic. To evaluate the performance of our approach, we compiled a list of CMs from the literature. Since all these CMs have either been confirmed experimentally or have been identified by at least three published studies we shall assume that they are high-confidence CMs (Supplementary Table S1). We shall use this compiled reference list to help guide our choice of p-value cut-off by considering the true positive rate (TPR, Supplementary Figure S2).

We were able to recover 84.6 % (15/17, Supplementary Table S1) of published high-confidence CMs using a heuristic p-value cut-off set at the 98 % interval, resulting in a preliminary list of mutations that are significantly associated with RIF resistance. Synonymous mutations are unlikely to have a strong effect on the growth phenotype, as they do not change the structure of the final protein and thus they were removed from the list. The resulting list contained 78 putative CMs (Supplementary Table S2).

We mapped these 78 putative CMs onto a phylogenetic tree of the *M. tuberculosis* samples (Methods). The majority of the most frequent putative CMs in our dataset indeed show homoplasy (Figure 3A). Interestingly, the putative CM E1092D on *rpoC* exclusively occurred in one single clade of Lineage 2 and is thus highly likely to be a phylogenetic marker rather than a CM. We observed clustering in clades of Lineage 2 for other CMs (Figure 3A: P1040R, I491V: blue trapezoid and V483A: red trapezoid), but in contrast to E1092D, these CMs also occurred in other, genetically distant parts of the tree. This makes convergent evolution of compensation more likely than homology.

**Figure 3:**
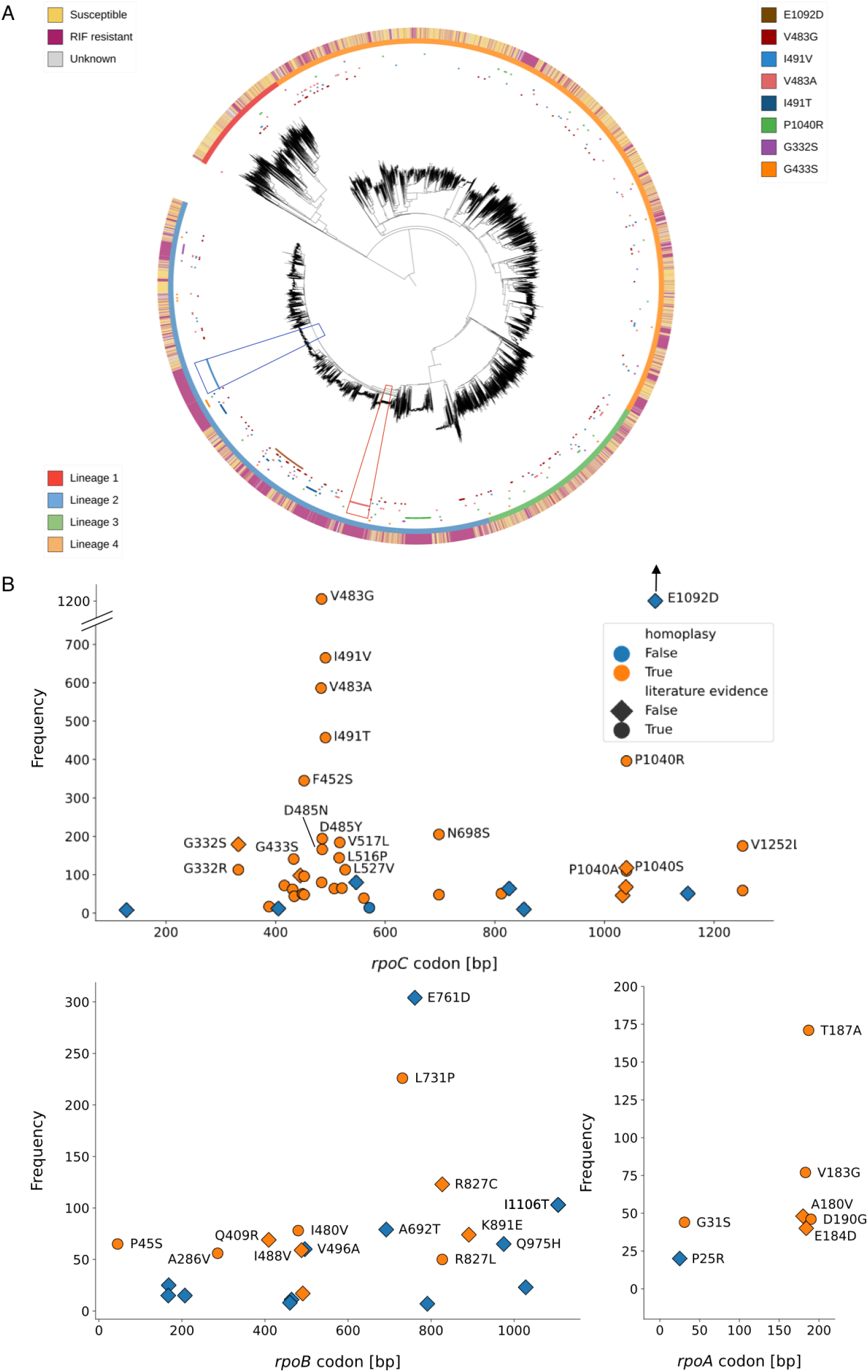
Putative CMs are distributed widely across the phylogenetic tree and the RNA polymerase genes. **(A)** Phylogenetic tree assembled from single nuceotide polymorphism (SNP)-distances of about 15,000 samples. The resistance level of the samples is indicated on the outermost ring, while the second ring indicates the lineage. The most common CMs were mapped on the innermost ring, with the trapezoids indicating clades within Lineage 2 that show a cluster of a specific CM (I491V: blue trapezoid and V483A: red trapezoid). **(B)** The putative CMs were mapped according to their position in the respective gene and their frequency of co-occurrence with resistance. There were five hits on the *σ* factor that are not shown, as well as one hit on the *rpoZ* gene (Supplementary Table S2). All of these do not show homoplasy. E1092D is outside of the plotting range due to its high frequency (1989 observations).

After filtering out hits that do not exhibit homoplasy, we arrived at a final list of 51 putative CMs. Out of these 51 hits, 12 have to our knowledge not been previously described (Figure 3B, Supplementary Table S2). All these hits would be valid starting points for further investigations of compensatory effects and mechanisms.

### Most compensatory mutations are found at interfaces between the RNA polymerase subunits

To evaluate their position on the protein structure, we mapped all 51 non-synonymous putative CMs that show homoplasy (Supplementary Table S2) onto the RNAP in complex with RIF. ^13^ The hits clustered in four different regions of the RNAP (Figure 4A): i) at the interfaces between subunits (Figure 4B), ii) on the *β* and *β ^′^* subunits close to the “rifampicin resistance determining region” (RRDR) and the active center (Figure 4C), iii) around the secondary channel in the *β ^′^* subunit (Figure 4D) and iv) at the DNA entry channel (Figure 4E). We found putative CMs in the *β* subunit of the RNAP, which had long been overlooked as a possible location for CMs, as most efforts in identifying CMs initially focused on the *β ^′^* subunit. ^30^ There were putative CMs in most subunits of the RNAP, however only the ones in the *rpoA*, *rpoB* and *rpoC* genes show homoplasy (Figure 3B, Supplementary Table S2). Still, most of the homoplastic CMs were found on the *β ^′^* subunit, which supports the initial hypothesis that it plays the major part in compensation.

**Figure 4:**
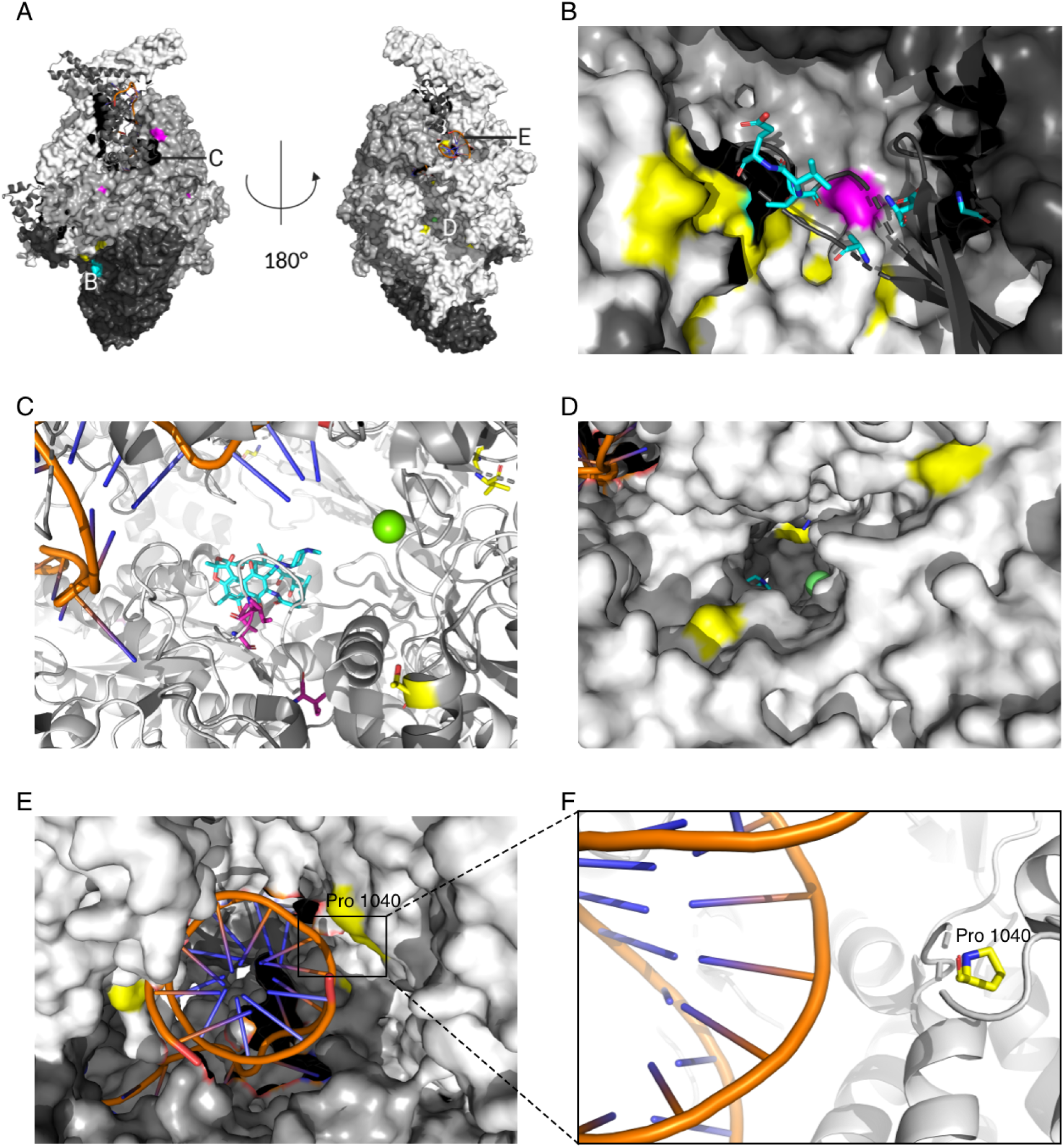
Compensatory mutations (CMs) map to various subunits of the RNA polymerase (RNAP) **(A)** Overview of clustering regions for CMs. Letters indicate where the CM clustering regions are located. **(B)** The interaction region of subunits *α* (black), *β* (dark grey) and *β ^′^* (light grey). CMs can be found in all these subunits and are highlighted in color (*β* subunit: magenta, *β ^′^* subunit: yellow, *α* subunit: light blue, stick representation). **(C)** CMs close to the ‘rifampicin resistance determining region’ (RRDR) on the *β* and *β ^′^* subunits. Rifampicin (RIF, light blue) is shown bound to the RRDR. The DNA strand is visible on the top left in dark blue and orange stick representation. The active center magnesium ion is shown in green. CMs are highlighted in colour (*β* subunit: magenta and *β ^′^* subunit: yellow) and by stick representation. **(D)** CMs close to the RNAP secondary channel in the *β ^′^* subunit (light grey) are shown in yellow. The location of the active site inside the protein can be deduced through the active site magnesium ion, indicated in green. **(E)** CMs close to the DNA entry channel are shown in yellow. The DNA helix is shown in dark blue and orange stick representation. **(F)** Close-up of the location of a putative CM (yellow stick representation, CM mutates Proline to Arginine) close to the DNA backbone. This CM might change interactions of the RNAP with the DNA strand.

The majority of CMs, including those most common in our dataset (Supplementary Table S2: V483G/A and I491V/T), were located close to the interface of the *β*, *β ^′^* and *α* RNAP subunits (Figure 4B, Supplementary Figure S3). CMs in the interfacial region might alter binding of the subunits, perhaps leading to higher protein stability, without necessarily affecting the active site. ^31^ Several hits also clustered close to the RRDR, where RIF binds to the RNAP of susceptible bacteria (Figure 4C). Mutations close to the RRDR have been suspected to alter the conformation of the active site, possibly leading to altered activity of the enzyme. ^30^

The RNAP secondary channel (Figure 4D) and the DNA entry channel (Figure 4E) are locations for CMs that, to our knowledge, have not been discussed in the literature. The secondary channel serves as a direct connection from the outside of the protein to the active center and is presumed to facilitate the diffusion of substrate nucleotides into the protein for incorporation into the nascent mRNA. It has been proposed that molecules entering through this channel could regulate RNAP activity as well. ^37^ CMs at this location could therefore modify diffusion in and out of the RNAP. Mutated residues in the DNA entry channel could alter interactions with the DNA helix. This location is especially interesting, since we observed CMs very close to the DNA strand where it enters the protein (Figure 4F).

### Rifampicin resistant *M. tuberculosis* samples show higher *in vitro* growth densities in presence of compensatory mutations in Lineages 2 and 3

If we have correctly identified CMs, growth densities in the resistant samples with CMs should be higher than in resistant samples without CMs. To test this we compared the growth distributions of pan- susceptible samples, RIF resistant samples that contain at least one homoplastic CM from our complete list (Supplementary Table S2), and resistant samples without any CMs.

Resistant samples with CMs grew to significantly higher densities (p = 3.92e-57) than resistant samples without CMs (Figure 5A, Supplementary Table S3). Surprisingly, the median growth of resistant samples with CMs also significantly out-performed the growth of pan-susceptible samples (p = 6.25e-26, Figure 5A, Supplementary Table S4). The association of CMs with such high growth levels could mean that CMs increase fitness above wild-type levels, or hint at confounding factors.

**Figure 5:**
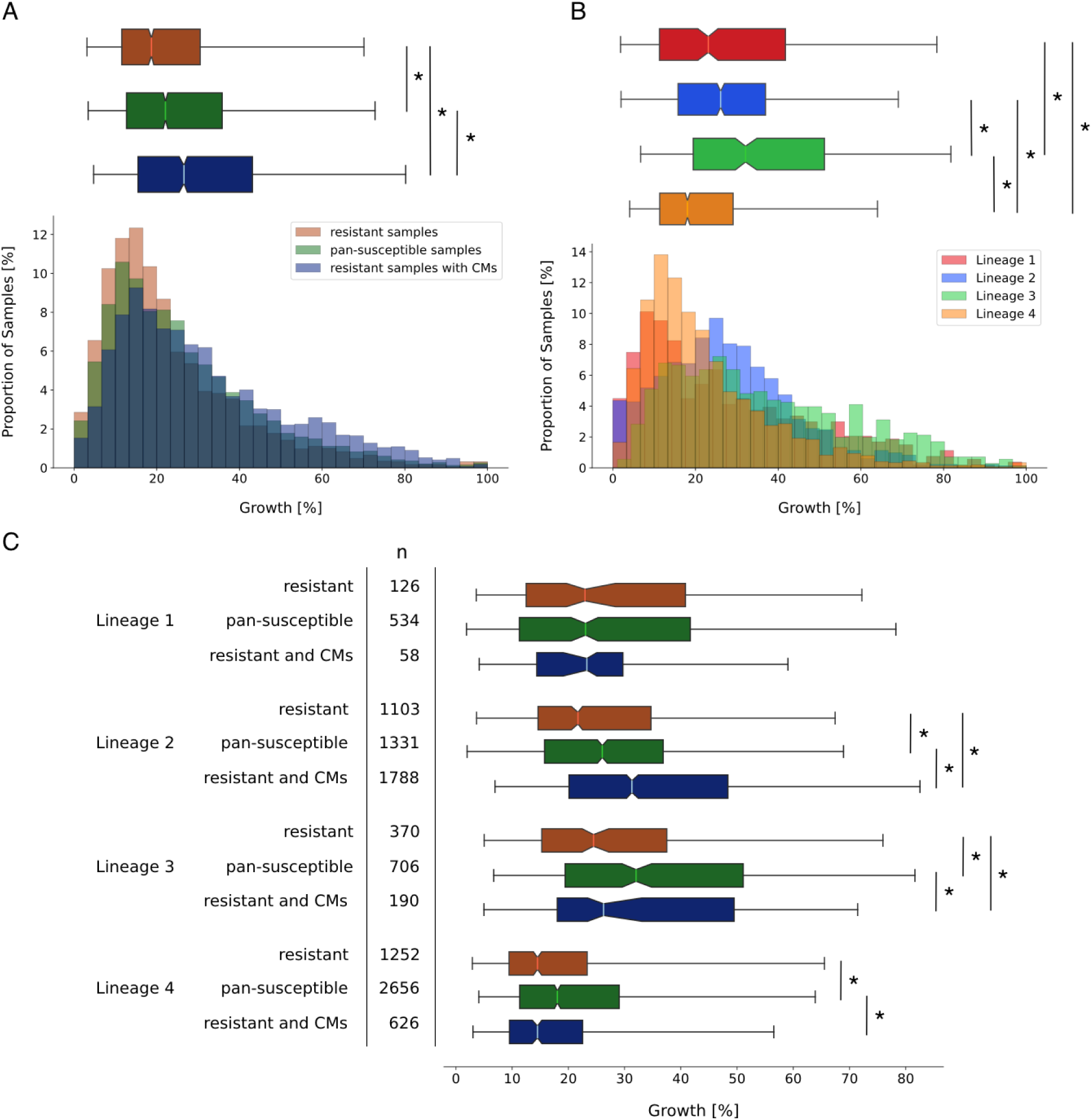
Presence of compensatory mutations (CMs) in samples with rifampicin (RIF) resistance- conferring mutations in the RNA polymerase of *M. tuberculosis* is associated with higher growth densities in some lineages. **(A)** Distributions of growth in percent of covered well-area as measured in the CRyPTIC project ^33^ were plotted as a histogram against the proportion of samples that display this amount of growth (bottom) and as a notched box plot reflecting the distribution quantiles (top). Samples with RIF resistance mutations but no putative CMs are plotted in red, samples that were classified as pan-susceptible are plotted in green. Samples that have RIF resistance mutations and at least one CM are shown in blue. For the box plot, half of the data lies within the area of the box and 95% in the area covered by the whiskers. Outliers (5 % of the data) were removed to achieve a cleaner representation. Indented area close to the medians indicate their respective confidence interval, while the star (*) indicates a significant Mann-Whitney p-value (p *<* 0.05 %). The respective medians, confidence intervals and the Mann-Whitney p-values are listed in Supplementary Table S3.**(B)** Plot structure equivalent to the box plot in A, but the bars represent pan-susceptible samples from different lineages, plotted in the respective colours indicated in the legend. The respective medians, confidence intervals and the Mann-Whitney p-value are listed in Supplementary Table S4. **(C)** Plot structure equivalent to the box plot in A, but the box plots represent subsets of samples that belong to the lineage displayed on the left. The sample size is shown in the column marked with ‘n’. For a histogram representation of the same data refer to Supplementary Figure S4. The respective medians, confidence intervals and the Mann-Whitney p-values are listed in Supplementary Table S5.

One confounding factor could be rooted in the differences in virulence between *M. tuberculosis* lineages. Lineage 2 is thought to be associated with higher transmission rates than other lineages ^38^, which could be reflected in higher *in vitro* growth densities. Indeed, considering only pan-susceptible samples, Lineage 2 showed significantly higher growth than Lineage 4 (p = 7.99e-33) and higher growth than Lineage 1 (p = 7.17e-02), whilst still being out-performed by Lineage 3 (p = 8.36e-16, Figure 5B, Supplementary Table S4). In addition, 60% of resistant samples in Lineage 2 contained at least one CM from our final list, compared to 34% in other lineages. Higher growth densities of Lineage 2 combined with CM accumulation could lead to inflated growth phenotype association with CMs.

We hence need to examine lineage contribution to the enhanced growth phenotype in CM samples. In Lineage 2 we still found a significantly higher growth density for samples with CMs than for pansusceptible samples (p = 3.23e-02, Figure 5C). For all other lineages, the growth difference between resistant samples with CMs and pan-susceptible samples was either insignificant, or pan-susceptible samples grew to higher densities (Figure 5C, Supplementary Table S5). In Lineage 3 we observed a significantly higher growth density of resistant samples with CMs when compared to resistant samples without CMs (p = 2.08e-02). This indicates that while CMs partially restore fitness, the extreme growth phenotypes are not caused by CMs alone. There might be other growth-associated mutations in Lineage 2, acting as confounding factors. We do not observe an effect on growth in Lineages 1 and 4. In Lineage 1, the sample size might be too low to detect an effect on growth, since we could not even observe the initial fitness cost in resistant samples without CMs when compared to pan-susceptible samples (Figure 5C, Supplementary Table S5). Lineage 4 on the other hand showed very low overall growth (Figure 5B) which makes it difficult to detect the positive effect of CMs due to the relatively small initial effect of resistance mutations on growth (Figure 5C).

### The effect of compensatory mutations on *in vitro* growth is confounded with lineage and clade affiliation

Lineage 2 is the only lineage that showed significantly higher growth in resistant samples with CMs than in pan-susceptible samples and it also accumulates CMs more than any other lineage. While this might indicate that CMs play an important role in this Lineage, we suspected that there might be other mutations co-occurring with CMs that enhanced growth densities to the reported high levels.

We hence wanted to investigate if the high growth densities observed in samples with CMs within Lineage 2 reflect the growth advantage of a few high-fitness clades. We aimed to dissect the clade influence for two representative CM clusters with sufficient available samples (Figure 3A). The first cluster showed accumulation of the CM I491V (blue trapezoid in Figure 3A). The average SNP-distance of 23 within the cluster indicated that this cluster is not the result of recent transmission according to UKHSA guidelines ^39^, although the maximum SNP-distance of 44 highlights that samples are nonetheless very closely related. Most likely the cluster is indeed an established clade within Lineage 2 and not an oversampled local outbreak. We saw that the clade with the CM I491V cluster showed significantly higher growth than pan-susceptible samples (p = 6.52e-24), but the effect outside of this clade was just as strong (p = 1.12e-07, Figure 6A, Table 2). It is however possible that there are other Lineage 2 clades associated with this CM that confounded the increased growth. The second cluster showed accumulation of the CM V483A (red trapezoid in Figure 3A). The average SNP-distance was higher than in the other cluster, with around 31 pairwise SNP differences between samples in the cluster and a maximum SNP-distance of 149. We saw that median growth in the clade with the CM V483A cluster was significantly higher than in samples with CMs outside of the clade (p = 6.79e-03). But median growth of the latter was still significantly higher than the median growth of resistant samples without CMs (p = 4.24e-03, Figure 6B, Table 2), which suggests that a small positive effect of CMs on growth is conserved after removing the confounding influence of the high-fitness clade.

**Figure 6:**
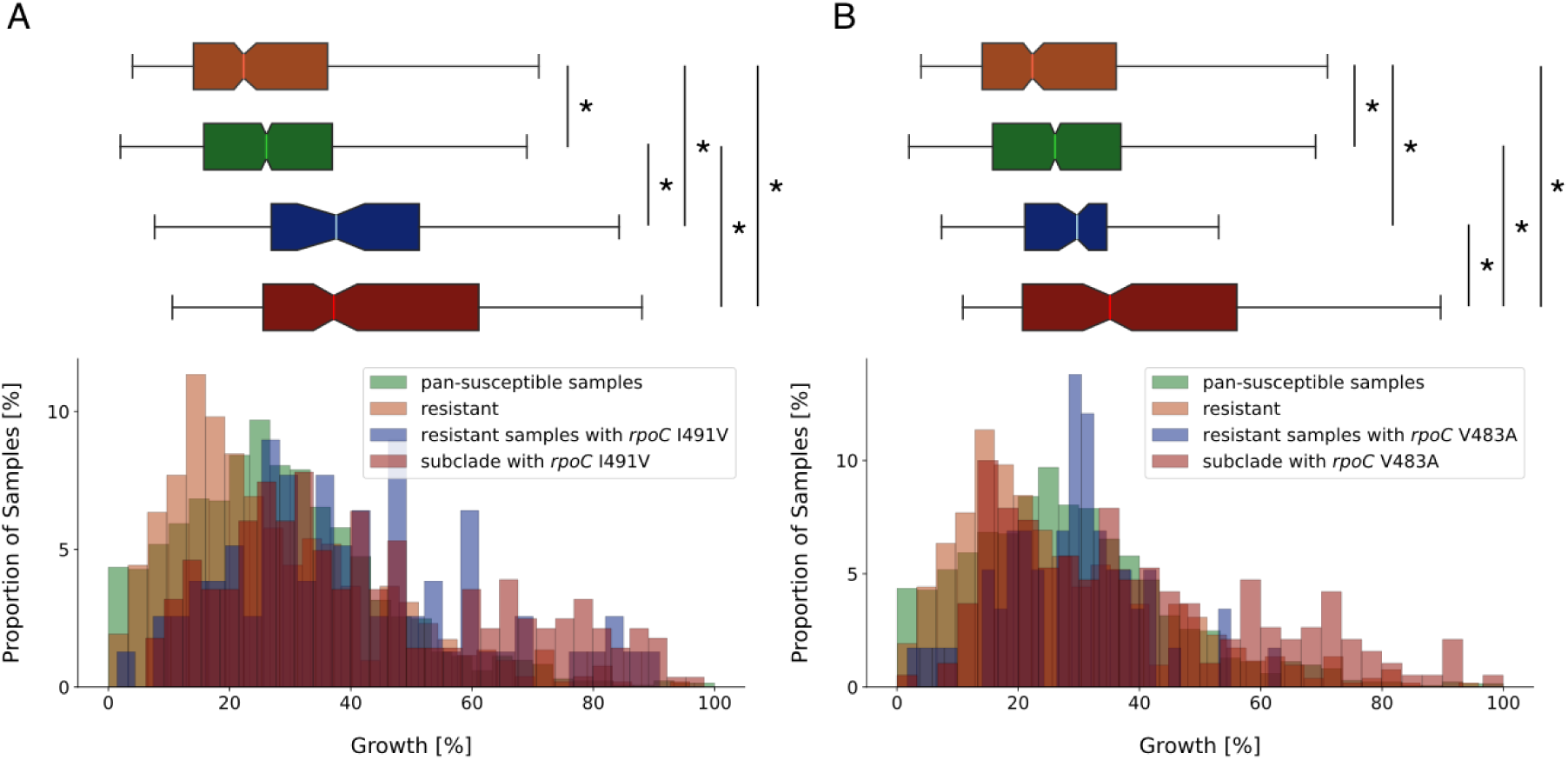
*M. tuberculosis* clades with clusters of compensatory mutations (CMs) explain some of the high growth densities associated with CMs in Lineage 2. **(A)** Growth distributions (percentage of covered well area) in Lineage 2 were plotted as a histogram against the proportion of samples that display this amount of growth (bottom) and as a notched box plot reflecting the distribution quantiles (top). Lineage 2 samples were classified as pan-susceptible (green), RIF resistant (red), resistant and showing the CM I491V outside of the CM cluster clade (blue) and as part of the Lineage 2 clade where all samples show the CM I491V (dark red). For the box plot, half of the data lies within the area of the box and 95% in the area covered by the whiskers. Outliers (5% of the data) were removed to achieve a cleaner representation. Indented area close to the medians indicate their respective confidence interval, while the star (*) indicates a significant Mann-Whitney p-value (p *<* 0.05%). The respective medians, confidence intervals and the Mann-Whitney p-values are listed in Table 2. **(B)** Plot structure equivalent to the box plot in A, but the CM in question is V483A. Distribution medians and their confidence intervals are shown in Table 2.

**Table 2:** Median growth of samples from Lineage 2 with different compensatory mutations, compared to growth of pan-susceptible samples, growth of resistant samples without CMs and growth of a resistant clade that shows a cluster of the respective CM. The confidence interval (CI) for the median is calculated using bootstrapping where ‘CI low’ indicates the lower threshold and ‘CI high’ the upper threshold. P-values are given with respect to resistant sample growth (p-value*_r_*), pan-susceptible sample growth (p-value*_s_*) and sample growth of resistant samples with CMs outside of the cluster subclade (p-value*_rCM_*). n indicates the sample size.

| sample type | median growth [%] | CI low | CI high | p-value<sub>r</sub> | p-value<sub>s</sub> | p-value<sub>rCM</sub> | n |
| --- | --- | --- | --- | --- | --- | --- | --- |
| pan-susceptible | 26.08 | 25.21 | 26.99 |  |  |  | 1331 |
| resistant and no CMs | 21.6 | 20.5 | 22.4 |  | 2.46e-05 |  | 1103 |
| CM *rpoC* I491V: |  |  |  |  |  |  |  |
| resistant | 37.6 | 31.5 | 42.3 | 9.57e-11 | 1.12e-07 |  | 78 |
| resistant and subclade | 37.2 | 33.9 | 41.4 | 2.14e-33 | 6.52e-24 | 0.499 | 282 |
| CM *rpoC* V483A: |  |  |  |  |  |  |  |
| resistant | 29.7 | 26.9 | 31.7 | 4.24e-03 | 0.263 |  | 58 |
| resistant and subclade | 35.1 | 30.7 | 38.9 | 9.45e-17 | 9.89e-11 | 6.79e-03 | 190 |

The association of CM clusters with specific high-fitness clades within Lineage 2 hence does appear to be one confounding factor in our growth data analysis and might explain the observed increase of growth densities to levels higher than the wild-type growth densities. However, the small growth advantage of resistant sample with CMs over resistant sample without CMs appears to be conserved even when the high-fitness clades are removed from the analysis. Further work is needed to completely disentangle the influence of these high-fitness clades on the growth phenotype.

## Discussion

Resistance to first-line antimicrobial drugs like rifampicin (RIF) is wide-spread in *M. tuberculosis*, the etiological agent of tuberculosis (TB). Accordingly, investigating the emergence and spread of resistance is a matter of high interest in a world where antibiotics are one of our main defences against bacterial infections. A key factor in the spread of RIF resistance are compensatory mutations (CMs), named due to their potential to compensate for the fitness cost that resistance mutations introduce in *M. tuberculosis*.

In this study, we have identified a comprehensive set of high-confidence CMs in *M. tuberculosis* and also captured and dissected their influence on *in vitro* growth of the bacteria. Overall, we identified 78 unique high-confidence putative CMs, of which 51 exhibit homoplasy (Supplementary Table S2). This set includes 84.6 % of the compiled reference CMs (Supplementary Table S1) and 12 hits that are, to our knowledge, novel. Through mapping these hits on the RNAP structure, we identified four CM clustering regions (Figure 4A). Two of them have, to our knowledge, not been described before: the DNA entry tunnel and the RNAP secondary channel.

The fitness cost of RIF resistance is reflected in the growth distributions of samples with resistance versus samples with no resistance mutations (Figure 2A). Specifically, we observed that the reduced fitness in samples with resistance mutations is recapitulated by our *in vitro* growth data. When considering the overall dataset, the presence of CMs appeared to not only restore the growth of resistant *M. tuberculosis* samples but also samples with CMs even exceeded the growth densities of pan-susceptible bacteria (Figure 5A). Since we are assuming growth is a proxy for fitness, this would suggest that strains with CMs are generally associated with higher fitness than pan-susceptible strains. However, when considering lineage affiliation, only Lineage 2 showed these unusually high growth in samples with CMs (Figure 5C), although Lineage 3 resistant samples with CMs still showed significantly higher growth than resistant samples (Supplementary Table S5). The absence of an effect in other lineages can be explained by lower sample sizes or lower overall growth.

The high growth densities in Lineage 2 appear to be caused by CM clusters associated with clades with increased growth phenotypes (Figure 6B), such as for *rpoC* V483A. While resistant samples with this CM still grew to higher densities than resistant samples with no CMs, they did not reach susceptible *in vitro* growth densities outside of the high-growth clade. This growth data analysis also made it clear that such investigations will have to be conducted lineage- or even sublineage-wise, to account for population structure. Concerning the impact of CMs on growth, our findings indicate that while CMs increase *in vitro* growth to a certain extent, they could be associated with and/or may act synergistically with other growth-enhancing factors that boost growth yet further, such as clade-specific mutations.

Their strong association with high-fitness clades makes CMs interesting candidates for further studies.

There are two possible scenarios, firstly it is possible that CMs may have been preceded by evolutionary older secondary growth-associated mutations that push growth above wild-type levels. This is supported by the fact that the high-growth CM clusters were located in sublineages 2.2 and 2.2.7, respectively, which are mostly part of the modern Beijing sublineages. These lineages are known to exhibit increased virulence ^40^. Secondly, it is possible that by partially restoring fitness after resistance acquisition, CMs could have facilitated the emergence of further mutations and hence the establishment of these high-fitness clades. This would make CMs a key determinant in the increased transmissibility observed in Lineage 2. In both scenarios, the notion that CMs play an important role in the spread of the increasingly virulent Lineage 2 is supported by the fact that this lineage accumulates CMs more than any other lineage, presumably due to the reported higher mutation rates ^41^. It has been suggested before that such compensated multi-drug resistant (MDR) mutants might be selected for in environments where many patients are treated with antituberculosis drugs ^40^. This emphasises the need for appropriate antibiotic stewardship to prevent further spread of MDR *M. tuberculosis*.

The conformational changes caused by CMs mostly affect the interfaces between the subunits and the space around the active site. These mutations are likely to enhance transcriptional activity of the RNAP either through changing the conformation of the active site or through increasing the overall stability of the protein complex. The increased activity could, in turn, lead to increased transmissibility. Whether CMs are directly associated with transmission rates is an ongoing discussion in the field ^42–46^, with most sources supporting the notion of CMs being associated with increased transmission. Our results support a positive influence of CMs on growth and, consequently, on fitness in samples with CMs in Lineages 2 and 3. This could translate to higher transmission rates at population level, although this has to be tested through *in vivo* experiments.

The nature of our growth data is a general limitation of our approach to capturing the fitness effects of CMs. Because growth was measured after two weeks of incubation, any information on fitness advantages visible during the exponential growth phase, such as growth rate, will be lost. This could be changed by including growth monitoring during the exponential growth phase of the bacteria. Since this is difficult for a slow-growing species like *M. tuberculosis*, polymerase activity assays could potentially close this gap. By testing laboratory-derived *M. tuberculosis* mutants with an engineered combination of resistance mutations and CMs, the direct influence of CMs on RNAP activity could be tested. Even so, as the growth data was acquired *in vitro*, any fitness advantages that arise from the interaction of *M. tuberculosis* bacteria with their host environment will not be captured. We hence cannot necessarily directly translate the results of this study to the epidemiological reality of *M. tuberculosis* spread in human populations. As this was outside the scope of this study, we also did not account for possible associations between mutations outside of the RNA polymerase (RNAP) and RIF resistance. One could however adapt our statistical association testing method for other genes.

Overall, the whole-genome sequencing data enabled the construction of a tractable, high-confidence list of 51 putative CMs, which could form the basis of future investigations. In addition, we derived further insights by combining the sequencing data with our growth data, allowing us to estimate the changes of the growth phenotype following emergence of resistance and CMs. CMs arise in a similar fashion in other *M. tuberculosis* genes, such as *ahpC* and *gyrA*, following resistance to isoniazid and fluoroquinolones, respectively. ^47^ In other bacterial species, such as *Salmonella typhimurium* and *Escherichia coli*, there is evidence for a fitness cost due to antibiotic resistance and compensatory mechanisms arising to reduce its impact. ^48^ Our approach to identifying CMs could hence be applied in a similar fashion to other microorganisms and drugs where resistance mutations are annotated and known to introduce a fitness cost.

## Acknowledgments

This work was supported by funding from the Biotechnology and Biological Sciences Research Council (UKRI-BBSRC) [grant number BB/T008784/1], by the National Institute for Health Research (NIHR) Oxford Biomedical Research Centre (BRC) and also by the National Institute for Health Research (NIHR) Health Protection Research Unit in Healthcare Associated Infections and Antimicrobial Resistance (NIHR200915), a partnership between the UK Health Security Agency (UKHSA) and the NIHR Biomedical Research Centre, Oxford. We are grateful for discussions with Profs Tim EA Peto, David Eyre, Daniel Wilson and Dr Kerri Malone. The views expressed are those of the authors and not necessarily those of the NHS, the NIHR, the Department of Health or the UK Health Security Agency. The funders had no role in study design, data collection and analysis, decision to publish, or preparation of the manuscript.

## Materials and Methods

### Reproducibility statement

The majority of figures and tables in this report can be reproduced using a GitHub repository and attendant jupyter notebook available online. ^49^

### Dataset sources

The initial dataset comprised 77,860 whole-genome sequenced patient-derived samples of *M. tuberculosis* ^32^ collected and collated by the CRyPTIC project. ^32^ Mutations with respect to version 3 of the H37Rv reference genome were aggregated and are available in a series of data tables, along with other data, such as the predicted resistances to various antibiotics, lineage association and the amount of growth in the 96-well plates. ^34^

The amount of bacterial growth in the two positive controls of each 96-well plate after two weeks incubation was obtained from photographs of each sample plate by some image processing software, AMyGDA. ^33^ Since this is only possible for CRyPTIC samples that were collected and subsequently plated, we only have growth data for 15,211 samples.

### Pairwise statistical association testing

We compiled a list of putative compensatory mutations (CMs) by performing statistical association testing for all mutations in the RNA polymerase (RNAP) genes that were observed to co-occur with any mutations listed as conferring resistance to rifampicin (RIF) in the catalogue. ^34^ We excluded any samples where sequencing was inconclusive. The statistical test used was Fisher’s exact test, which has a high computational cost that increases with dataset size. To minimise the effect of this, we used a Python3 package with an optimised implementation of Fisher’s test. This is faster than the standard implementation (**0** fishers-exact-test) and enabled us to run Fisher’s exact test for every pair of resistance and co-occurring mutation within 2.5 hours on a modern workstation.

### Evaluating and filtering results from Fisher’s exact test

Despite being generally accepted as a conservative approach to correct for multiple testing, naively applying a Bonferroni-corrected p-value of 0.01 % (Fig. S2) to the results of the pairwise Fisher’s exact test leads to hundreds of hits and a TPR of 100%, most likely due to the inflating effect of linkage disequilibrium (LD) on p-values as mentioned in the Results section. We decided to only include hits with a p-value above the 98% percentile, since this still recovers 84.6 % of the high-confidence CMs and gives us a reasonable number of hits that we can process using our downstream filters. The entire list without p-value cut-off is available online. ^49^ To make sure we filter out any phylogenetic mutations, we removed any synonymous mutations and tested the remaining for homoplasy. Synonymous mutations tend to be phylogenetic markers since they do not change the protein structure and hence should not be subject to any selectional pressure. Homoplasy is an indicator of selective pressure that causes similar mutations to arise in genetically and evolutionary distinct populations. Our resulting preliminary list of CMs, including their respective associated resistance mutation is shown in Supplementary Table S2.

### Constructing binary homoplasy index

We mapped our putative hits on the published CRyPTIC phylogenetic tree ^32^ using the publicly available software iTOL ^50^. Any samples originating from organisms other than *M. tuberculosis* (e.g. *Mycobacterium caprae* and *Mycobacterium bovis*) and lineages other than 1-4 were removed. We also removed any samples where sequencing was inconclusive, since these were initially excluded from our statistical association testing. We inferred homoplasy if a mutation appeared in at least 3 genetically distant samples without common ancestors in the tree. This number was chosen to partly account for the possibility of false positives due to sequencing errors or hemiplasies. It should be noted that if a mutation was not classified as homoplastic, this does not exclude the possibility of this mutation being a CM.

### Plotting of growth data

The average growth, as measured by AMyGDA ^33^ of the two positive control wells for each sample was obtained from the CRyPTIC dataset. To quantify the differences between growth of samples that are pan-susceptible, resistant or resistant with CMs, the pairwise Mann-Whitney p-value for the two growth distributions was calculated. In addition, the medians and confidence intervals of the medians were approximated using a bootstrapping approach and included in notched box-plot figures. This is a robust method considering the non-normal distribution of the growth data. For significance, we always referred to the Mann-Whitney p-values.

### Mapping of putative compensatory mutations on RNA polymerase structure

High-confidence homoplastic hits were mapped in PyMOL onto the crystal structure of the *M. tuberculosis* RNAP in complex with RIF ^13^ to identify clusters and elucidate potential compensatory mechanisms.

## References

1. H. S. Gold and R. C. Moellering (1996) New England Journal of Medicine 335:1445–1445.

2. H. Nikaido (2009) Annual Review of Biochemistry 78:119–146.

3. K. S. Ikuta, L. R. Swetschinski, G. R. Aguilar, F. Sharara, T. Mestrovic, A. P. Gray, N. D. Weaver, E. E. Wool, C. Han, A. G. Hayoon, et al. (2022) The Lancet 400:2221–2221.

4. P. A. Smith and F. E. Romesberg (2007) Nature Chemical Biology 3:549–549.

5. K. Iskandar, J. Murugaiyan, D. Hammoudi Halat, S. E. Hage, V. Chibabhai, S. Adukkadukkam, C. Roques, L. Molinier, P. Salameh, and M. Van Dongen (2022) Antibiotics 11:182.

6. E. Tacconelli, E. Carrara, A. Savoldi, S. Harbarth, M. Mendelson, D. L. Monnet, C. Pulcini, G. Kahlmeter, J. Kluytmans, Y. Carmeli, et al. (2018) The Lancet Infectious Diseases 18:318–318.

7. P. E. Almeida Da Silva and J. C. Palomino (2011) Journal of Antimicrobial Chemotherapy 66:1417–1417.

8. E. Cambau and M. Drancourt (2014) Clinical Microbiology and Infection 20:196–196.

9. S. Bagcchi (2023) The Lancet Microbe 4:e20.

10. E. D. Chan and M. D. Iseman (2008) Current Opinion in Infectious Diseases 21:587–587.

11. M. Merker, C. Blin, S. Mona, N. Duforet-Frebourg, S. Lecher, E. Willery, M. G. B. Blum, S. Rüsch-Gerdes, I. Mokrousov, E. Aleksic, et al. (2015) Nature Genetics 47:242–242.

12. A. Bortoluzzi, F. W. Muskett, L. C. Waters, P. W. Addis, B. Rieck, T. Munder, S. Schleier, F. Forti, D. Ghisotti, M. D. Carr, et al. (2013) J Biol Chem 288:14438–14438.

13. W. Lin, S. Mandal, D. Degen, Y. Liu, Y. W. Ebright, S. Li, Y. Feng, Y. Zhang, S. Mandal, Y. Jiang, et al. (2017) Molecular Cell 66:169–179.e8.

14. S. M. Gygli, S. Borrell, A. Trauner, and S. Gagneux (2017) FEMS Microbiology Reviews 41:354–354.

15. L. K. W. Yuen, D. Leslie, and P. J. Coloe (1999) Journal of Clinical Microbiology 37:3844–3844.

16. P. A. Aristoff, G. A. Garcia, P. D. Kirchhoff, and H. D. Hollis Showalter (2010) Tuberculosis 90:94–94.

17. D. M. Rothstein (2016) Cold Spring Harbor Perspectives in Medicine 6:a027011.

18. O. J. Billington, T. D. McHugh, and S. H. Gillespie (1999) Antimicrobial Agents and Chemotherapy 43:1866–1866.

19. S. Gagneux, C. D. Long, P. M. Small, T. Van, G. K. Schoolnik, and B. J. M. Bohannan (2006) Science 312:1944–1944.

20. M. A. Stefan, F. S. Ugur, and G. A. Garcia (2018) Antimicrob Agents Chemother 62:e00164–18.

21. S. Maisnier-Patin and D. I. Andersson (2004) Research in Microbiology 155:360–360.

22. G. Brandis, M. Wrande, L. Liljas, and D. Hughes (2012) Molecular Microbiology 85:142–142.

23. A. K. Alame Emane, X. Guo, H. E. Takiff, and S. Liu (2021) Tuberculosis 129:102091.

24. T. Song, Y. Park, I. C. Shamputa, S. Seo, S. Y. Lee, H.-S. Jeon, H. Choi, M. Lee, R. J. Glynne, S. W. Barnes, et al. (2014) Molecular Microbiology 91:1106–1106.

25. I. Comas, S. Borrell, A. Roetzer, G. Rose, B. Malla, M. Kato-Maeda, J. Galagan, S. Niemann, and S. Gagneux (2012) Nature Genetics 44:106–106.

26. M. de Vos, B. Müller, S. Borrell, P. A. Black, P. D. van Helden, R. M. Warren, S. Gagneux, and T. C. Victor (2013) Antimicrobial Agents and Chemotherapy 57:827–827.

27. Q.-j. Li, W.-w. Jiao, Q.-q. Yin, F. Xu, J.-q. Li, L. Sun, J. Xiao, Y.-j. Li, I. Mokrousov, H.-r. Huang, et al. Antimicrobial Agents and Chemotherapy 60:2807–2812.

28. N. Casali, V. Nikolayevskyy, Y. Balabanova, O. Ignatyeva, I. Kontsevaya, S. R. Harris, S. D. Bentley, J. Parkhill, S. Nejentsev, S. E. Hoffner, et al. (2012) Genome Research 22:735–735.

29. A. Ali, Z. Hasan, R. McNerney, K. Mallard, G. Hill-Cawthorne, F. Coll, M. Nair, A. Pain, T. G. Clark, and R. Hasan (2015) PLOS ONE 10:e0117771.

30. P. Ma, T. Luo, L. Ge, Z. Chen, X. Wang, R. Zhao, W. Liao, and L. Bao (2021) Emerging Microbes & Infections 10:743–743.

31. V. Ruiz and A. Paula (2020) Int J Mycobacteriol 9:121–121.

32. The CRyPTIC Consortium (2022) PLoS Biology 20:e3001721. https://ftp.ebi.ac.uk/pub/databases/cryptic/releasejune2022/.

33. P. W. Fowler, A. L. Gibertoni Cruz, S. J. Hoosdally, L. Jarrett, E. Borroni, M. Chiacchiaretta, P. Rathod, S. Lehmann, N. Molodtsov, T. M. Walker, et al. (2018) Microbiology 164:1522–1522.

34. The CRyPTIC Consortium and the 100,000 Genomes Project, C. Allix-Béguec, I. Arandjelovic, L. Bi, P. Beckert, M. Bonnet, P. Bradley, A. M. Cabibbe, I. Cancino-Muñoz, M. J. Caulfield, et al. (2018) The New England Journal of Medicine 379:1403–1415.

35. S. G. Earle, C.-H. Wu, J. Charlesworth, N. Stoesser, N. C. Gordon, T. M. Walker, C. C. A. Spencer, Z. Iqbal, D. A. Clifton, K. L. Hopkins, et al. (2016) Nature Microbiology 1:1–1.

36. P. E. Chen and B. J. Shapiro (2021) Classic genome-wide association methods are unlikely to identify causal variants in strongly clonal microbial populations. preprint, Genomics.

37. B. E. Nickels and A. Hochschild (2004) Cell 118:281–281.

38. M. Karmakar, J. M. Trauer, D. B. Ascher, and J. T. Denholm (2019) Journal of Infection 79:572–581.

39. UK Health Security Agency (2022). Mycobacterium tuberculosis whole-genome sequencing and cluster investigation handbook.

40. S. C. M. Ribeiro, L. L. Gomes, E. P. Amaral, M. R. M. Andrade, F. M. Almeida, A. L. Rezende, V. R. Lanes, E. C. Q. Carvalho, P. N. Suffys, I. Mokrousov, et al. (2014) Journal of Clinical Microbiology 52:2615–2615.

41. C. B. Ford, R. R. Shah, M. K. Maeda, S. Gagneux, M. B. Murray, T. Cohen, J. C. Johnston, J. Gardy, M. Lipsitch, and S. M. Fortune (2013) Nat Genet 45:784–784.

42. S. M. Gygli, C. Loiseau, L. Jugheli, N. Adamia, A. Trauner, M. Reinhard, A. Ross, S. Borrell, R. Aspindzelashvili, N. Maghradze, et al. (2021) Nature Medicine 27:1171–1171.

43. S. Wang, Y. Zhou, B. Zhao, X. Ou, H. Xia, Y. Zheng, Y. Song, Q. Cheng, X. Wang, and Y. Zhao (2020) Front Med 14:51–51.

44. M. Merker, M. Barbier, H. Cox, J.-P. Rasigade, S. Feuerriegel, T. A. Kohl, R. Diel, S. Borrell, S. Gagneux, V. Nikolayevskyy, et al. eLife 7:e38200.

45. Y. Chen, Q. Liu, H. E. Takiff, and Q. Gao (2022) Journal of Infection 85:49–49.

46. Q. Liu, T. Zuo, P. Xu, Q. Jiang, J. Wu, M. Gan, C. Yang, R. Prakash, G. Zhu, H. E. Takiff, et al. (2018) Emerg Microbes Infect 7:98.

47. G. Napier, S. Campino, J. E. Phelan, and T. G. Clark (2023) Sci Rep 13:623.

48. M. G. Reynolds (2000) Genetics 156:1471–1471.

49. V. Brunner and P. W. Fowler. Accompanying code to reproduce analysis and figures. https://github.com/fowler-lab/tb-rnap-compensation.

50. I. Letunic and P. Bork (2021) Nucleic Acids Research 49:W293–W296.

