## Supplemental Information for "Compensatory mutations are associated with increased *in vitro* growth in resistant clinical samples of *Mycobacterium tuberculosis*"

| putative CM | Exp. evidence | 1 | 2 | 3 | 4 | 5 | 6 | 7 | 8 | sum(ref) | fisher |
|---|---|---|---|---|---|---|---|---|---|---|---|
| *rpoC* V483G | ✓ | ✓ | ✓ | ✓ | ✓ | ✓ | ✓ | ✓ | | 7 | ✓ |
| *rpoC* V483A | ✓ | ✓ | ✓ | ✓ | ✓ | | | ✓ | | 5 | ✓ |
| *rpoC* I491T | | ✓ | ✓ | ✓ | | ✓ | ✓ | | | 5 | ✓ |
| *rpoC* L527V | | ✓ | ✓ | | ✓ | ✓ | | | | 4 | ✓ |
| *rpoA* T187A | | ✓ | ✓ | | | ✓ | | | ✓ | 4 | ✓ |
| *rpoC* G332R | | | | ✓ | ✓ | ✓ | | ✓ | | 4 | ✓ |
| *rpoC* L507V | | | ✓ | | ✓ | | ✓ | ✓ | | 4 | |
| *rpoC* N698S | | ✓ | ✓ | ✓ | ✓ | | | | | 4 | ✓ |
| *rpoC* W484G | | ✓ | ✓ | | | | | | ✓ | 3 | ✓ |
| *rpoC* I491V | ✓ | ✓ | ✓ | | ✓ | | | | | 3 | ✓ |
| *rpoC* V517L | | ✓ | ✓ | | ✓ | | | | | 3 | ✓ |
| *rpoC* L516P | ✓ | ✓ | | | ✓ | | | ✓ | | 3 | ✓ |
| *rpoC* H525Q | | | ✓ | ✓ | ✓ | | | | | 3 | |
| *rpoC* N698K | ✓ | ✓ | | | ✓ | | ✓ | | | 3 | ✓ |
| *rpoC* V1252L | ✓ | | | | ✓ | | ✓ | ✓ | | 3 | ✓ |
| *rpoC* F452L | ✓ | | | | | | ✓ | | | 1 | ✓ |
| *rpoC* P1040R | ✓ | | | | ✓ | | | | | 1 | ✓ |

**Table S1:** Reference CMs used for evaluating our approach to identifying new CMs. Putative CMs are shown on the left. We included reference CMs that were either proven experimentally or identified in at least three of the reference papers. Fisher indicates if the CM came up as significantly resistance associated in our statistical association test and was located below the heuristic p-value threshold and showed homoplasy.

| resistance | putative CM | only CM | both | -$\log_{10}$(p-value) | literature evidence | homoplasy |
|---|---|---|---|---|---|---|
| *rpoB* S450L | *rpoC* E1092D | 2012 | 1989 | inf | | |
| *rpoB* S450L | *rpoC* V483G | 37 | 1206 | inf | ✓ | ✓ |
| *rpoB* S450L | *rpoC* I491V | 19 | 665 | inf | ✓ | ✓ |
| *rpoB* S450L | *rpoC* V483A | 33 | 586 | inf | ✓ | ✓ |
| *rpoB* S450L | *rpoC* I491T | 10 | 457 | 293.06 | ✓ | ✓ |
| *rpoB* S450L | *rpoC* P1040R | 32 | 396 | 225.09 | ✓ | ✓ |
| *rpoB* S450L | *rpoC* F452S | 2 | 345 | 230.58 | ✓ | ✓ |
| *rpoB* S450L | *rpoB* E761D | 0 | 304 | 207.05 | | |
| *rpoB* S450L | *rpoB* L731P | 1 | 226 | 151.44 | ✓ | ✓ |
| *rpoB* S450L | *rpoC* N698S | 2 | 205 | 135.23 | ✓ | ✓ |
| *rpoB* S450L | *rpoC* D485Y | 8 | 194 | 118.89 | ✓ | ✓ |
| *rpoB* S450L | *rpoC* V517L | 1 | 184 | 122.87 | ✓ | ✓ |
| *rpoB* S450L | *rpoC* G332S | 16 | 179 | 100.19 | | ✓ |
| *rpoB* S450L | *rpoC* V1252L | 3 | 175 | 113.24 | ✓ | ✓ |
| *rpoB* S450L | *rpoA* T187A | 3 | 171 | 110.54 | ✓ | ✓ |
| *rpoB* S450L | *rpoC* D485N | 2 | 166 | 108.82 | ✓ | ✓ |
| *rpoB* S450L | *rpoC* L516P | 3 | 144 | 92.37 | ✓ | ✓ |
| *rpoB* S450L | *rpoC* G433S | 3 | 141 | 90.35 | ✓ | ✓ |
| *rpoB* S450L | *rpoB* R827C | 8 | 123 | 72.06 | | ✓ |
| *rpoB* S450L | *rpoC* P1040S | 3 | 118 | 74.93 | | ✓ |
| *rpoB* S450L | *rpoC* L527V | 3 | 113 | 71.58 | ✓ | ✓ |
| *rpoB* S450L | *rpoC* G332R | 5 | 113 | 68.95 | ✓ | ✓ |
| *rpoB* S450L | *rpoC* P1040A | 1 | 110 | 72.70 | ✓ | ✓ |
| *rpoB* L452P | *rpoB* I1106T | 3 | 103 | 212.49 | | |
| *rpoB* D435G | *rpoB* I1106T | 3 | 103 | 263.70 | | |
| *rpoB* S450L | *rpoC* K445R | 0 | 98 | 66.49 | | ✓ |
| *rpoB* S450L | *rpoC* F452L | 3 | 96 | 60.24 | ✓ | ✓ |
| *rpoB* S450L | *rpoC* L547V | 2 | 80 | 50.94 | | |
| *rpoB* S450L | *rpoC* W484G | 11 | 80 | 41.69 | ✓ | ✓ |
| *rpoB* S450L | *rpoB* A692T | 16 | 79 | 37.45 | | |
| *rpoB* S450L | *rpoB* I480V | 3 | 78 | 48.27 | ✓ | ✓ |
| *rpoB* S450L | *rpoA* V183G | 10 | 77 | 40.62 | ✓ | ✓ |
| *rpoB* S450L | *rpoB* K891E | 0 | 74 | 50.18 | | ✓ |
| *rpoB* S450L | *rpoC* N416S | 3 | 72 | 44.30 | ✓ | ✓ |
| *rpoB* S450L | *rpoB* Q409R | 14 | 69 | 32.77 | | ✓ |
| *rpoB* S450L | *rpoC* V1039A | 1 | 68 | 44.37 | | ✓ |
| *rpoB* S450L | *rpoB* P45S | 5 | 65 | 37.49 | ✓ | ✓ |
| *rpoB* S450L | *rpoB* Q975H | 17 | 65 | 28.57 | | |
| *rpoB* S450L | *rpoC* A521D | 2 | 65 | 40.93 | ✓ | ✓ |
| *rpoB* S450L | *rpoC* L507V | 1 | 64 | 41.68 | ✓ | ✓ |
| *rpoB* S450L | *rpoC* N826T | 0 | 64 | 43.39 | | |
| *rpoB* S450L | *rpoC* V431M | 4 | 62 | 36.58 | ✓ | ✓ |
| *rpoB* S450L | rpoB V496A | 0 | 60 | 40.68 | | |
| *rpoB* S450L | *rpoB* I488V | 2 | 59 | 36.94 | | ✓ |
| *rpoB* S450L | rpoC V1252M | 2 | 59 | 36.94 | ✓ | ✓ |
| *rpoB* S450L | *rpoB* A286V | 4 | 56 | 32.68 | ✓ | ✓ |
| *rpoB* S450L | *rpoC* T812I | 3 | 51 | 30.48 | ✓ | ✓ |
| rpoB S450L | *rpoC* K1152Q | 0 | 51 | 34.57 | | |
| *rpoB* S450L | *rpoB* R827L | 0 | 50 | 33.89 | ✓ | ✓ |
| *rpoB* S450L | *rpoC* L449V | 0 | 50 | 33.89 | ✓ | ✓ |

| | | | | | | |
|---|---|---|---|---|---|---|
| *rpoB* S450L | *rpoC* N698K | 4 | 48 | 27.50 | $\checkmark$ | $\checkmark$ |
| *rpoB* S450L | *rpoC* F452C | 1 | 48 | 30.94 | $\checkmark$ | $\checkmark$ |
| *rpoB* S450L | *rpoA* A180V | 0 | 48 | 32.54 | | $\checkmark$ |
| *rpoB* I491F | *rpoC* E1033A | 25 | 46 | 95.60 | | $\checkmark$ |
| *rpoB* S450L | *rpoA* D190G | 0 | 46 | 31.18 | $\checkmark$ | $\checkmark$ |
| *rpoB* S450L | *rpoA* G31S | 0 | 44 | 29.82 | $\checkmark$ | $\checkmark$ |
| *rpoB* S450L | *rpoC* P434R | 3 | 44 | 25.91 | $\checkmark$ | $\checkmark$ |
| *rpoB* S450L | *rpoA* E184D | 0 | 40 | 27.11 | | $\checkmark$ |
| *rpoB* H445R | *rpoC* S561P | 2 | 39 | 98.27 | $\checkmark$ | $\checkmark$ |
| *rpoB* V170F | *rpoB* V168A | 2 | 25 | 64.98 | | |
| *rpoB* L452P | *rpoB* H1028R | 1 | 23 | 46.47 | | |
| *rpoB* S450W | *rpoA* P25R | 1 | 20 | 45.52 | | |
| *rpoB* H445D | *rpoC* G388A | 1 | 17 | 32.33 | $\checkmark$ | $\checkmark$ |
| *rpoB* D435G | *rpoB* I491L | 18 | 17 | 32.89 | | $\checkmark$ |
| *rpoB* D435Y | *rpoB* R167C | 1 | 15 | 29.72 | | |
| *rpoB* S450W | *sigA* A223T | 0 | 15 | 35.07 | | |
| *rpoB* H445Y | *rpoB* E207K | 0 | 15 | 29.05 | | |
| *rpoB* V170F | *rpoC* G571R | 11 | 14 | 30.91 | $\checkmark$ | |
| *rpoB* S441A | *rpoC* L405M | 28 | 12 | 36.05 | | |
| *rpoB* S441A | *rpoB* L464M | 3 | 11 | 39.12 | | |
| *rpoB* Q432P | *rpoC* T853A | 3 | 10 | 28.72 | | |
| *rpoB* Q432K | *rpoZ* T107I | 1 | 10 | 30.13 | | |
| *rpoB* S441A | *sigA* E385Q | 14 | 9 | 27.77 | | |
| *rpoB* S441A | *sigA* G380A | 3 | 9 | 31.34 | | |
| *rpoB* S441A | *sigA* I382V | 3 | 9 | 31.34 | | |
| *rpoB* S441A | *sigA* L386M | 4 | 9 | 30.83 | | |
| *rpoB* S441A | *rpoB* E460D | 5 | 8 | 26.66 | | |
| *rpoB* S441A | *rpoC* I128V | 3 | 8 | 27.56 | | |
| *rpoB* S441A | *rpoB* R791T | 0 | 7 | 25.92 | | |

**Table S2:** Hit list resulting from Fisher's exact test for association of resistance with co-occuring mutations, after removing synonymous mutations. The first column indicates the resistance mutation that the putative compensatory mutation (CM) in the second column is associated to. 'Only CM' indicates how often the CM occurs on its own, without the corresponding resistance mutation, and 'both' indicates how often we see the two mutations occur together. The last two columns indicate if the CM has been mentioned in the literature and if it shows homoplasy, respectively.

| sample type | median growth [%] | CI low | CI high | p-value$_r$ | p-value$_s$ | n |
|---|---|---|---|---|---|---|
| pan-susceptible | 22.1 | 21.6 | 22.7 | | | 5283 |
| resistant and no CMs | 18.7 | 18.0 | 19.3 | | 3.92e-15 | 2869 |
| resistant and CMs | 26.6 | 25.5 | 27.4 | 3.92e-57 | 6.25e-26 | 2667 |

**Table S3:** Median growth of resistant samples with compensatory mutations compared to pan-susceptible samples and samples with only resistance mutations. The confidence interval (CI) for the median is calculated using bootstrapping where 'CI low' indicates the lower threshold and 'CI high' the upper threshold. P-values are given with respect to resistant (p-value$_r$) and pan-susceptible sample growth (p-value$_s$) and n indicates the sample size.

| Lineage | median growth [%] | CI low | CI high | p-value$_1$ | p-value$_2$ | p-value$_3$ | n |
|---|---|---|---|---|---|---|---|
| Lineage 1 | 23.1 | 20.6 | 25.5 | | | | 534 |
| Lineage 2 | 26.1 | 25.3 | 27.0 | 7.17e-02 | | | 1331 |
| Lineage 3 | 32.1 | 29.6 | 34.8 | 8.68e-14 | 8.36e-16 | | 706 |
| Lineage 4 | 18.1 | 17.6 | 18.6 | 1.22e-05 | 7.99e-33 | 3.31e-64 | 2656 |

**Table S4:** Median growth of pan-susceptible samples from different *M. tuberculosis* lineages. The confidence interval (CI) for the median is calculated using bootstrapping where 'CI low' indicates the lower threshold and 'CI high' the upper threshold. P-values are given with respect to each lineage, indicated by the subscript x (p-value$_x$) and n indicates the sample size.

| sample type | median growth [%] | CI low | CI high | p-value$_r$ | p-value$_s$ | n |
|---|---|---|---|---|---|---|
| Lineage 1: | | | | | | |
| pan-susceptible | 23.1 | 20.6 | 25.3 | | | 534 |
| resistant and no CMs | 22.9 | 19.6 | 28.7 | | 0.784 | 126 |
| resistant and CMs | 23.3 | 19.6 | 25.2 | 0.516 | 0.596 | 58 |
| Lineage 2: | | | | | | |
| pan-susceptible | 26.1 | 25.4 | 27.2 | | | 1331 |
| resistant and no CMs | 21.6 | 20.4 | 22.5 | | 2.46e-05 | 1103 |
| resistant and CMs | 31.4 | 30.4 | 32.5 | 5.65e-43 | 3.20e-24 | 1788 |
| Lineage 3: | | | | | | |
| pan-susceptible | 32.1 | 29.6 | 34.9 | | | 706 |
| resistant and no CMs | 24.4 | 22.4 | 27.2 | | 8.79e-09 | 370 |
| resistant and CMs | 26.3 | 23.7 | 33.2 | 2.07e-02 | 4.38e-02 | 190 |
| Lineage 4: | | | | | | |
| pan-susceptible | 18.1 | 17.6 | 18.6 | | | 2656 |
| resistant and no CMs | 14.4 | 13.7 | 15.2 | | 6.17e-17 | 1252 |
| resistant and CMs | 14.6 | 13.7 | 15.4 | 0.995 | 3.44e-11 | 626 |

**Table S5:** Lineage-wise median growth of samples with different compensatory mutations compared to pan-susceptibles and samples with only resistance. The confidence interval (CI) for the median is calculated using bootstrapping where 'CI low' indicates the lower threshold and 'CI high' the upper threshold. P-values are given with respect to resistant (p-value$_r$) and pan-susceptible sample growth (p-value$_s$) and n indicates the sample size.

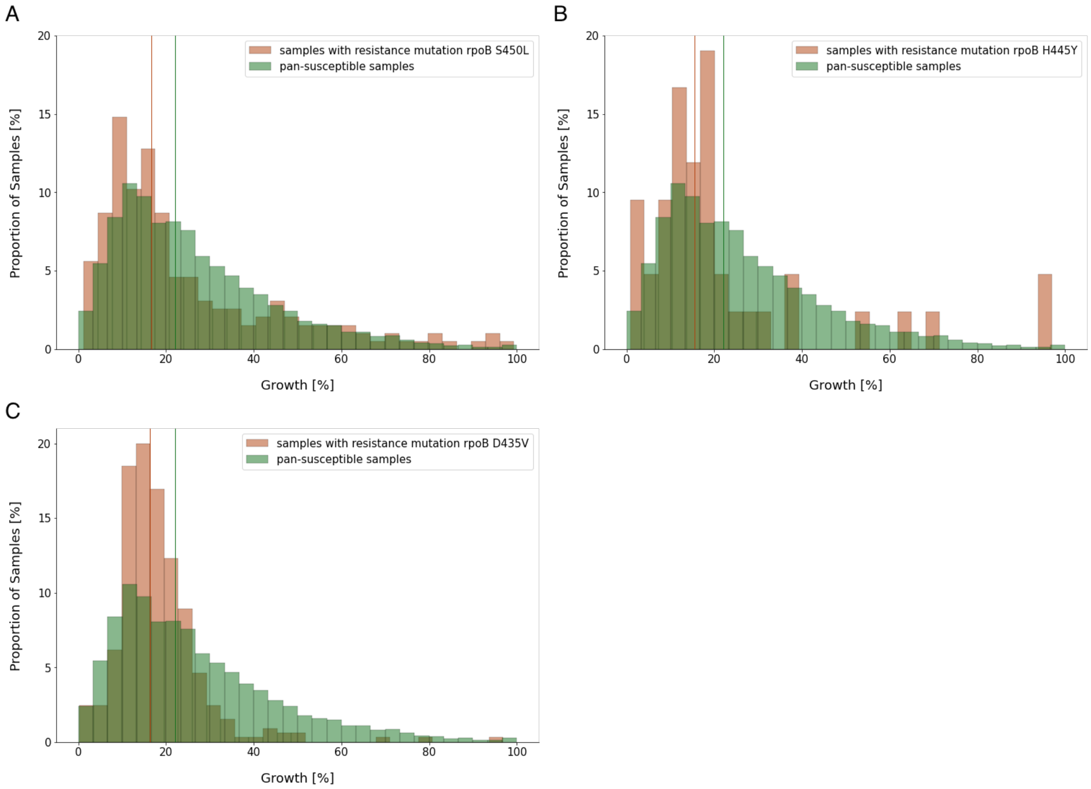


**Figure S1: Growth distributions for pan-susceptible samples vs samples with specific rifampicin (RIF) resistance mutations in *M. tuberculosis* (A-C)** Distributions of growth in percent of covered well-area as measured in the CRyPTIC project[9] were plotted as a histogram against the proportion of samples that display this amount of growth. Samples with the resistance mutation indicated in the legend and no other potentially interfering mutations are plotted in red, samples that were classified as pan-susceptible are plotted in green. Vertical lines indicate the respective medians. The medians and Mann-Whitney p-values of the distributions are listed in Table 1.

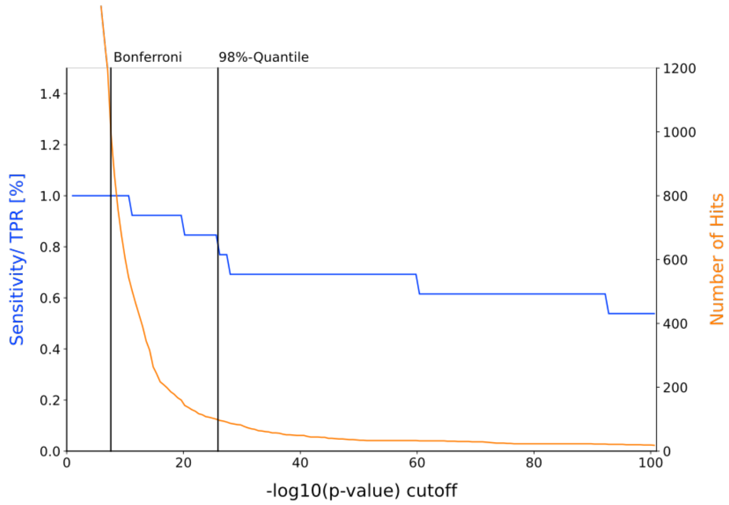


**Figure S2: Sensitivity and number of significant hits (putative compensatory mutations) depending on p-value** The graph shows the number of significant hits and reference hits detected depending on the $\log_{10}$ p-value cutoff shown on the x-axis. The left y-axis refers to the percentage of found reference hits from a compiled list, also termed sensitivity or true positive rate (TPR). The right y-axis shows the number of mutations that were classified as significantly resistance associated under the respective cut-off. The vertical lines indicate the p-value cut-off with Bonferroni correction and our heuristic p-value cut-off at the 98% quantile, respectively.

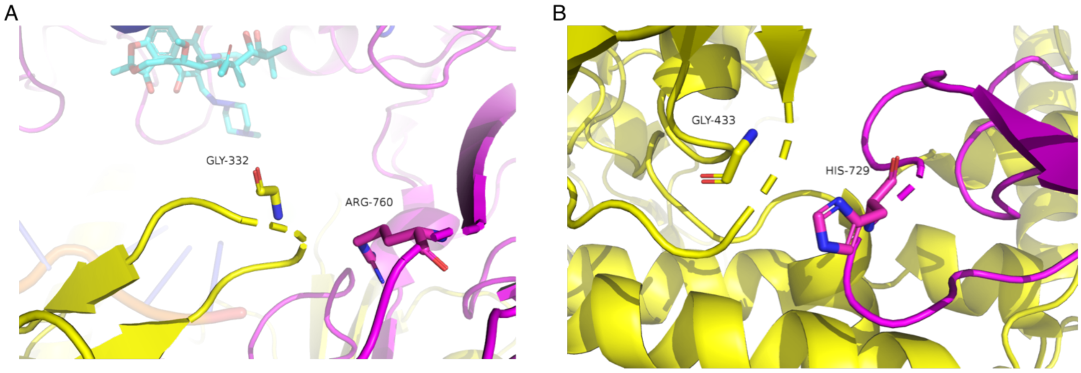


**Figure S3: Location of two high-confidence compensatory mutations (CMs) on the RNA polymerase (RNAP) (A)** The CM G332S is located on the $\beta'$ subunit, in a contact region to the $\beta$ subunit (magenta). The change from Glycine (stick representation) to Serine (negatively charged side chain) might enable an interaction with the close-by Arginine (positively charged side chain) on the $\beta$ subunit. The bound drug rifampicin (light blue) can be seen in the background. **(B)** The CM G433S is located on the $\beta'$ subunit, in a contact region to the $\beta$ subunit. The change from Glycine (stick representation) to Serine might enable an interaction with the close-by Histidine (positively charged side chain) on the $\beta$ subunit.

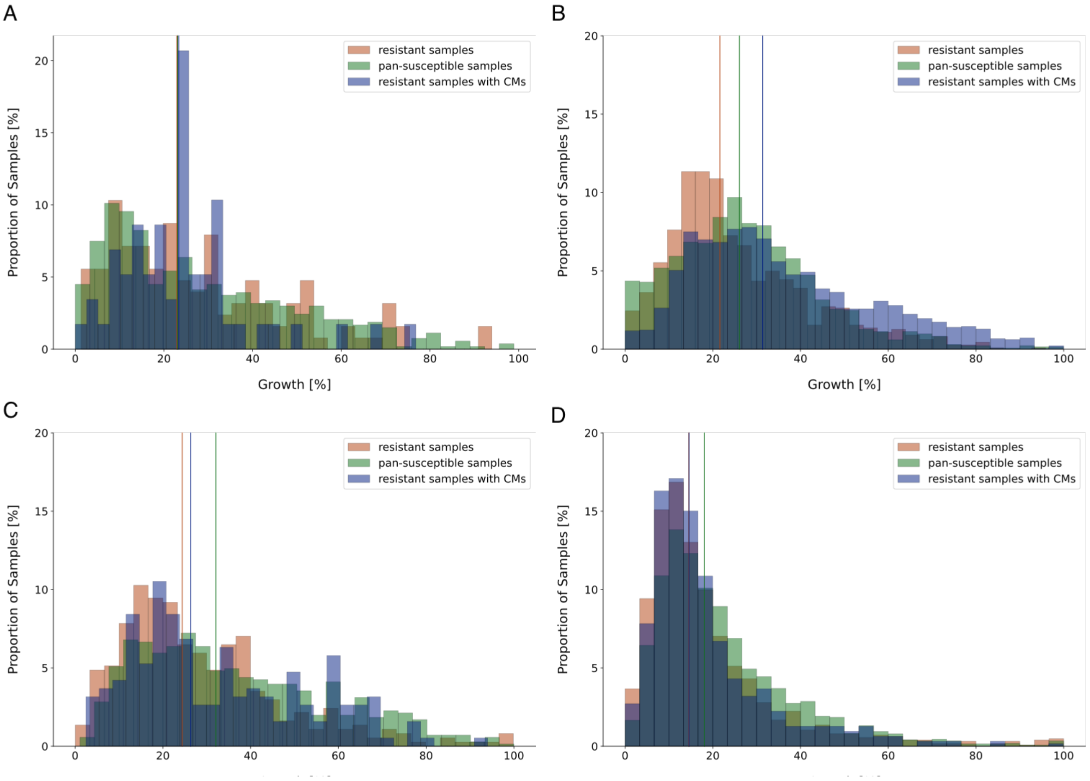


**Figure S4: Growth distributions of *M. tuberculosis* samples within different lineages (A)** Distribution of growth in *M. tuberculosis* Lineage 1 in percent of covered well-area as measured in the CRyPTIC project[9] were plotted as a histogram against the proportion of samples that display this amount of growth. Samples with rifampicin (RIF) resistance mutations but no putative compensatory mutations (CMs) are plotted in red, samples that were classified as pan-susceptible are plotted in green. Samples that have RIF resistance mutations and at least one CM are shown in blue. Vertical lines indicate the respective medians. The medians and Mann-Whitney p-values of the distributions are shown in Supplementary Table S5. **(B)** Plot layout as in (A), but samples derive from *M. tuberculosis* Lineage 2. **(C)** Plot layout as in (A), but samples derive from *M. tuberculosis* Lineage 3. **(D)** Plot layout as in (A), but samples derive from *M. tuberculosis* Lineage 4.

### References


[1] I. Comas, S. Borrell, A. Roetzer, G. Rose, B. Malla, M. Kato-Maeda, J. Galagan, S. Niemann, and S. Gagneux (2012) *Nature Genetics* 44:106–110.

[2] Q.-j. Li, W.-w. Jiao, Q.-q. Yin, F. Xu, J.-q. Li, L. Sun, J. Xiao, Y.-j. Li, I. Mokrousov, H.-r. Huang, et al. *Antimicrobial Agents and Chemotherapy* 60:2807–2812.

[3] M. de Vos, B. Müller, S. Borrell, P. A. Black, P. D. van Helden, R. M. Warren, S. Gagneux, and T. C. Victor (2013) *Antimicrobial Agents and Chemotherapy* 57:827–832.

[4] V. Ruiz and A. Paula (2020) *Int J Mycobacteriol* 9:121–137.

[5] N. Casali, V. Nikolayevskyy, Y. Balabanova, O. Ignatyeva, I. Kontsevaya, S. R. Harris, S. D. Bentley, J. Parkhill, S. Nejentsev, S. E. Hoffner, et al. (2012) *Genome Research* 22:735–745.

[6] T. Song, Y. Park, I. C. Shamputa, S. Seo, S. Y. Lee, H.-S. Jeon, H. Choi, M. Lee, R. J. Glynne, S. W. Barnes, et al. (2014) *Molecular Microbiology* 91:1106–1119.

[7] A. Ali, Z. Hasan, R. McNerney, K. Mallard, G. Hill-Cawthorne, F. Coll, M. Nair, A. Pain, T. G. Clark, and R. Hasan (2015) *PLOS ONE* 10:e0117771.

[8] P. Ma, T. Luo, L. Ge, Z. Chen, X. Wang, R. Zhao, W. Liao, and L. Bao (2021) *Emerging Microbes & Infections* 10:743–752.

[9] P. W. Fowler, A. L. Gibertoni Cruz, S. J. Hoosdally, L. Jarrett, E. Borroni, M. Chiacchiaretta, P. Rathod, S. Lehmann, N. Molodtsov, T. M. Walker, et al. (2018) *Microbiology* 164:1522–1530.
